# Dynamic influences on static measures of metacognition

**DOI:** 10.1101/2020.10.29.360453

**Authors:** Kobe Desender, Luc Vermeylen, Tom Verguts

## Abstract

Humans differ in their capability to judge the accuracy of their own choices via confidence judgments. Signal detection theory has been used to quantify the extent to which confidence tracks accuracy via M-ratio, often referred to as metacognitive efficiency. This measure, however, is static in that it does not consider the dynamics of decision making. This could be problematic because humans may shift their level of response caution to alter the tradeoff between speed and accuracy. Such shifts could induce unaccounted-for sources of variation in the assessment of metacognition. Instead, evidence accumulation frameworks consider decision making, including the computation of confidence, as a dynamic process unfolding over time. We draw on evidence accumulation frameworks to examine the influence of response caution on metacognition. Simulation results demonstrate that response caution has an influence on M-ratio. We then tested and confirmed that this was also the case in human participants who were explicitly instructed to either focus on speed or accuracy. We next demonstrated that this association between M-ratio and response caution was also present in an experiment without any reference towards speed. The latter finding was replicated in an independent dataset. In contrast, when data were analyzed with a novel dynamic measure of metacognition, which we refer to as v-ratio, in all of the three studies there was no effect of speed-accuracy tradeoff. These findings have important implications for research on metacognition, such as its measurement, domain-generality, individual differences, and neural correlates.

## Introduction

When asked to explicitly report how sure they are about their decisions, humans often claim high confidence for correct and low confidence for incorrect decisions. This capacity to evaluate the accuracy of decisions is often referred to as metacognitive accuracy. Although metacognitive accuracy about perceptual decisions is generally high ^1^, it varies significantly between participants ^2^ and between conditions ^3^. Such differences in metacognitive accuracy have important real-life consequences, as they relate, for example, to political extremism ^4^ and psychiatric symptoms ^5^. Moreover, an increasing number of researchers across different fields are starting to investigate to what extent observers can evaluate their own performance in different domains of cognition, such as sensorimotor uncertainty ^6^, motor movements and imagery ^7^, metric error monitoring ^8^, value-based decisions ^9^, group-decision making ^10^ and probabilistic learning^11^.

A debated question is how to quantify metacognitive accuracy. One prominent issue why one cannot simply calculate the correlation between confidence and choice accuracy ^12^ is that this confounds choice accuracy with metacognitive accuracy; i.e. it is much easier to detect one’s own mistakes in an easy task than in a hard task. Different solutions have been proposed in the literature, such as using coefficients from a logistic mixed-model ^13^, type 2 ROC curves ^2^, and meta-*d*’ ^14,15^. Rather than providing an in-depth discussion and comparison of these different measures, we here focus on one prominent static approach, namely the meta-*d*’ framework, the state-of-the-art measure of metacognitive accuracy ^16^. The meta-*d*’ approach is embedded within signal detection theory, and quantifies the extent to which confidence ratings discriminate between correct and incorrect responses (*meta-d*’) while controlling for first-order task performance (*d*’). Because both measures are on the same scale, one can calculate the ratio between both, meta-*d*’/*d*’, also called M-ratio, often referred to as metacognitive *efficiency*. When M-ratio is 1, all available first-order information is used in the (second-order) confidence judgment. When M-ratio is smaller than 1, metacognitive sensitivity is suboptimal, meaning that not all available information from the first-order response is used in the metacognitive judgment (Fleming & Lau, 2014). This measure has been used to address a variety of issues, such as whether metacognition is a domain-general capacity ^3,17,18^, the neural correlates of metacognition ^19–22^, how bilinguals differ from monolinguals ^23^, and how individual differences in metacognitive accuracy correlate with various constructs ^4,5^.

An important limitation is that the meta-*d*’ framework (just like the other static approaches cited above), does not consider dynamic aspects of decision making. Put simply, this measure depends on end-of-trial confidence and choice accuracy, but not on the response process governing the choice and its resulting reaction time. It is well known, however, that choice accuracy depends on response caution; i.e. choice accuracy decreases when participants are instructed to be fast rather than to be correct ^24,25^. The fact that static approaches of metacognition do not consider response caution is problematic because it confounds ability with caution: when focusing on speed rather than accuracy, one will produce many errors due to premature responding, and those errors are much easier to detect compared to errors resulting from low signal quality ^26^. Importantly, detecting “premature” errors does not imply “good metacognition” per se, but instead simply depends on one’s level of response caution.

To account for dynamic influences on metacognition, we propose to instead quantify metacognitive accuracy in a dynamic probabilistic framework ^27,28^. Sequential sampling models explain human decision making as a dynamic process of evidence accumulation ^29–31^. Specifically, decisions are conceptualized as resulting from the accumulation of noisy evidence towards one of two decision boundaries. The first boundary that is reached, triggers its associated decision. The height of the decision boundary controls the response caution with which a decision is taken ^24,25^. When lowering the boundary, decisions will be faster but less accurate; when increasing the boundary, decisions will be slower but more accurate. The prototypical dynamic sampling model is the drift diffusion model (DDM). In this model, confidence can be quantified as the probability of a choice being correct, given evidence, decision time, and the decision that was made ^32–34^ The relation between these three variables is represented by the heat map in Figure 1A. It captures the typical finding that trials with strong evidence are more likely to be correct than trials with weak evidence; and that trials with short RTs are more likely to be correct than trials with long RTs. As mentioned, the process of evidence accumulation terminates at the first boundary crossing. Formally, at that time the probability that the choice was correct can be quantified as *p*(*correct/e_t_, t, X*), where *e_t_* is the level of evidence at time *t, t* is the timing of boundary crossing and *X* is the choice made ^32,34,35^. In typical experiments, however, confidence judgments are provided separately in time (at time *t* + *s*, i.e., in a separate judgment after the choice), allowing evidence to further accumulate after boundary crossing. As a consequence, confidence should then be quantified as *p*(*correct/e_t+s_, t+s, X*), ^27,28,36^. This implies that a choice will be made once the integrated evidence reaches boundary *a*, but confidence is only computed after additional evidence accumulation (see Figure 1A for illustration).

**Figure 1.**
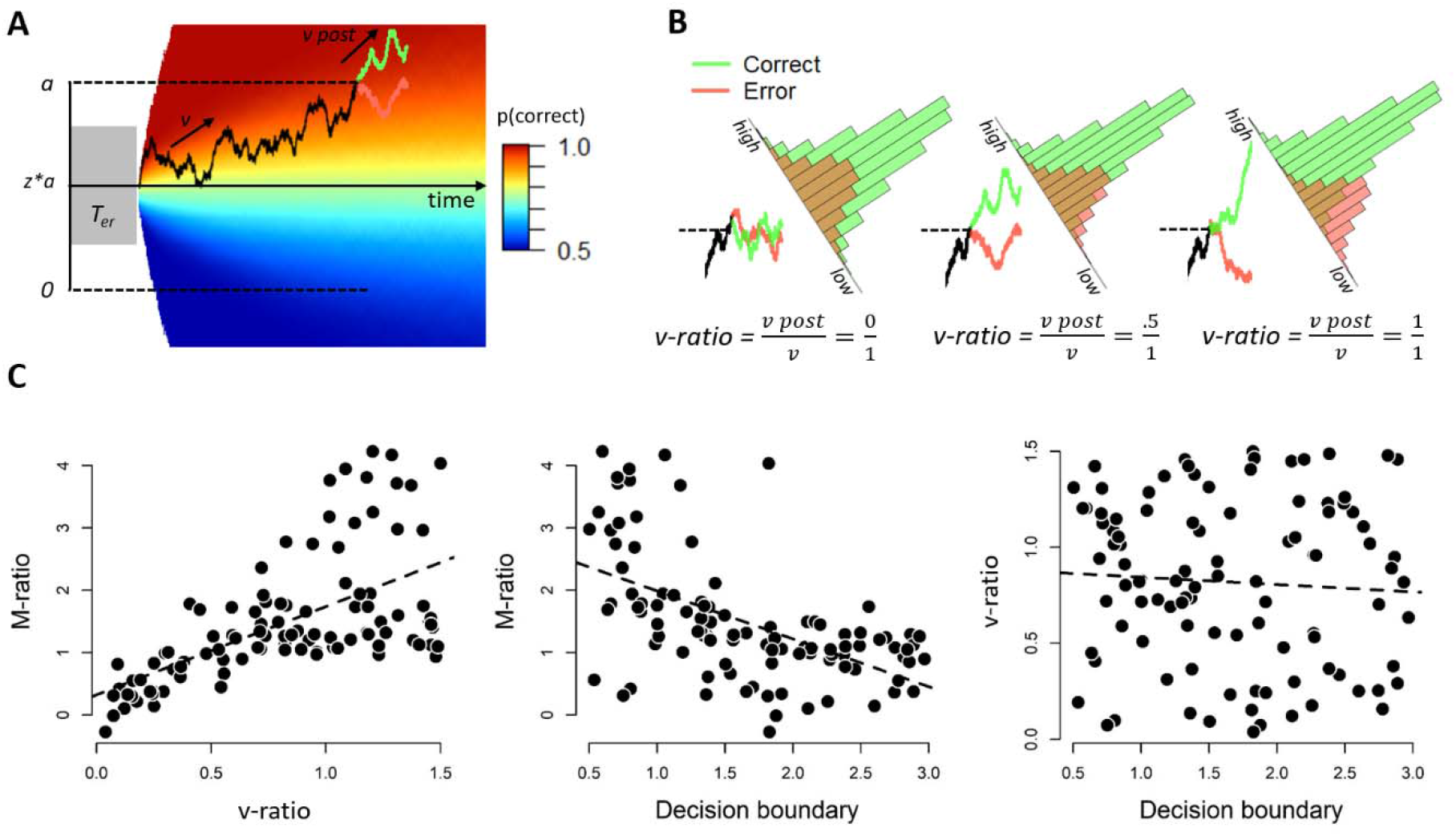
Quantifying metacognitive accuracy within an evidence accumulation framework. **A**. Noisy sensory evidence accumulates over time, until the integrated evidence reaches one of two decision boundaries (a or 0). After the decision boundary is reached, evidence continues to accumulate. The heat map (see Methods) shows the probability of being correct conditional on time, evidence, and the choice made (the choice corresponding to the upper boundary, in this example). Model confidence is quantified as this probability. **B**. Histograms of model-predicted confidence for different levels of v-ratio (reflecting the ratio between post-decision drift rate and drift rate). Higher levels of v-ratio are associated with better dissociating corrects from errors. **C.** Simulations from this dynamic evidence accumulation model show that M-ratio captures variation in v-ratio (r = .56; left panel), and critically, that M-ratio is also related to the differences in decision boundary (r = −.55; middle panel). By design, decision boundary and v-ratio are unrelated to each other (r ~ 0; right panel).

Within this formulation, good metacognitive accuracy can be considered as the ability to distinguish correct choices versus error choices based on *p*(*correct/e_t+s_, t+s, X*), i.e., based on confidence. Critically, the difference in the quantity *p*(*correct/e_t+s_, t+s, X*) for correct choices versus error choices, directly depends on the strength of post-decisional accumulation. This is visually depicted in Figure 1B: when post-decisional evidence accumulation is only driven by noise (i.e., post-decision drift rate is zero), model predicted confidence will be identical for correct and error trials (Figure 1B, left panel). On the other hand, when post-decisional evidence accumulation is very strong (i.e. post-decision drift rate is high) model predicted confidence strongly dissociates between corrects and errors, reflecting good metacognition (Figure 1B, right panel). From the above, it becomes clear that we can use the strength of post-decisional evidence accumulation as a dynamic measure of metacognitive accuracy. For comparison with the M-ratio framework, we quantified v-ratio as the ratio between post-decision drift rate and drift rate. Figure 1B shows post-decision accumulation for three scenarios with varying levels of v-ratio. As can be seen, if v-ratio is zero (left panel), additional evidence meanders adrift for both corrects and errors, and the model does not detect its own errors, i.e., representing a case of poor metacognitive accuracy. If, however, v-ratio equals 1 (i.e., post-decision drift rate and drift rate are the same), additional evidence confirms most of the correct choices (i.e., leading to high confidence) and disconfirms most of the error choices (i.e., leading to low confidence), i.e., representing good metacognitive accuracy. We thus propose that v-ratio can be used as a dynamic measure of metacognitive accuracy. In the following, we will shed light on the role of variation in response caution on both M-ratio and v-ratio.

## Results

### Model simulations reveal a link between response caution and M-ratio

We simulated data from a drift diffusion model with additional post-decisional evidence accumulation (see Figure 1A). Decision confidence was quantified as the probability of being correct given evidence, time and choice ^32,36,37^. We simulated data for 100 agents with 1000 observations each; for each agent, a different random value was selected for drift rate, non-decision time, decision boundary and post-decision drift rate. Importantly, we made sure that the simulated data showed the same correlation between choice RTs and confidence RTs as seen in the empirical data (see Methods). We then used these data to compute M-ratio, after dividing confidence ratings into four bins, separately for each observer (which is needed to compute meta-*d*’). As explained before, v-ratio was computed as the ratio between post-decision drift rate and drift rate. The results of our simulation study showed that, first, there was a clear positive relation between M-ratio and v-ratio, *r*(98) = .560, *p* < .001, reflecting that M-ratio captures individual variation in metacognition (Figure 1C, left panel). However, we also observed a strongly negative relation between M-ratio and decision boundary, *r*(98) = −.55, *p* < .001 (Figure 1C, central panel). This shows that M-ratio is highly dependent on the speed-accuracy tradeoff that one adopts: Lower bounds are associated with higher M-ratio. Intuitively, this occurs because lowering the decision boundary increases the probability of premature errors due to noise in the accumulation process. Given that M-ratio reflects a ratio between meta-*d*’ and *d*’, increasing the probability of premature errors can affect M-ratio in two ways: first, a lower decision boundary decreases *d*’, and will therefore have the effect that it increases M-ratio. Second, it is known that premature errors are easier to detect ^26^, and therefore a lower decision boundary might increase meta-*d*’, and will therefore increase M-ratio.

Finally, by design there was no relation between v-ratio and decision boundary, *r*(98) = −.065, *p* = .502 (Figure 1C, right panel). The full correlation matrix is shown in Table 1.

**Table 1.** Correlation table of the parameters from the model simulation. Note: ***<.001

|  | 1 | 2 | 3 | 4 | 5 |
| --- | --- | --- | --- | --- | --- |
| 1. Drift rate | - |  |  |  |  |
| 2. Non-decision time | -.21 | - |  |  |  |
| 3. Decision boundary | .13 | .11 | - |  |  |
| 4. v-ratio | .18 | -.03 | -.06 | - |  |
| 5. M-ratio | -.19 | .02 | -.55*** | .61*** | - |

### Experiment 1: Explicit speed-accuracy instructions affect static but not dynamic measures of confidence

Next, we tested these model predictions in an experiment with human participants. We recruited 36 human participants who performed a task that has been widely used in the study of evidence accumulation models: discrimination of the net motion direction in dynamic random dot displays ^29^. Participants were asked to decide whether a subset of dots was moving coherently towards the left or the right side of the screen (See Figure 2A). The percentage of dots that coherently moved towards the left or right side of the screen (controlling decision difficulty) was held constant throughout the experiment at 20%. After their choice, and a blank screen, participants indicated their level of confidence using a continuous slider. Critically, in each block, participants either received the instruction to focus on choice accuracy (“try to decide as accurate as possible”), or to focus on speed (“try to decide as fast as possible”). Consistent with the instructions, participants were faster in the speed condition than in the accuracy condition, *M_speed_* = 727ms versus *M_accuracy_* = 1014ms, *t*(35) = 4.47, *p* < .001, and numerically more accurate in the accuracy condition than in the speed condition, *M_accurate_* = 75.6% vs *M_speed_* = 73.8%, *t*(35) = 1.63, *p* = .111. Although there was no difference in overall accuracy, the speed-accuracy tradeoff manipulation introduced more “premature” errors in the speed condition compared to the accuracy condition (see Supplementary Figure 1). To test whether indeed errors in the speed condition were mostly “fast” errors and errors in the accuracy condition were mostly “slow” errors, we divided each participant’s error RTs into three equal-sized bins (fast, medium or slow). We observed that in the fast bin there were more errors from the speed than from the accuracy condition (*M^speed^* = 32.9 trials vs *M_accuracy_* = 14.4 trials, *t*(35) = −5.37,*p* < .001), whereas in the slow bin the reverse was true *(M_speed_* = 17.0 trials vs *M_accuracy_* = 31.0 trials, *t*(35) = 5.34,*p* < .001). In the medium bin the number of trials did not differ between both instruction conditions *(M_speed_* = 23.8 trials vs *M_accuracy_* = 23.8 trials, *p* = .974). Importantly, confidence RTs were not significantly different between the two instruction conditions, *p* = .249, suggesting participants selectively varied response caution for the motion coherence choice. The data further showed a significant between-participants correlation between median choice RTs and median confidence RTs in the accuracy condition, *r*(34) = .358, *p* = .032, but not in the speed condition, *r*(34) = .05, *p* = .778. Finally, participants were more confident in the accuracy condition than in the speed condition, *M_accuracy_* = 70 versus *M_speed_* = 67, *t*(35) = 3.57, *p* = .001 (See Figure 2D).

**Figure 2.**
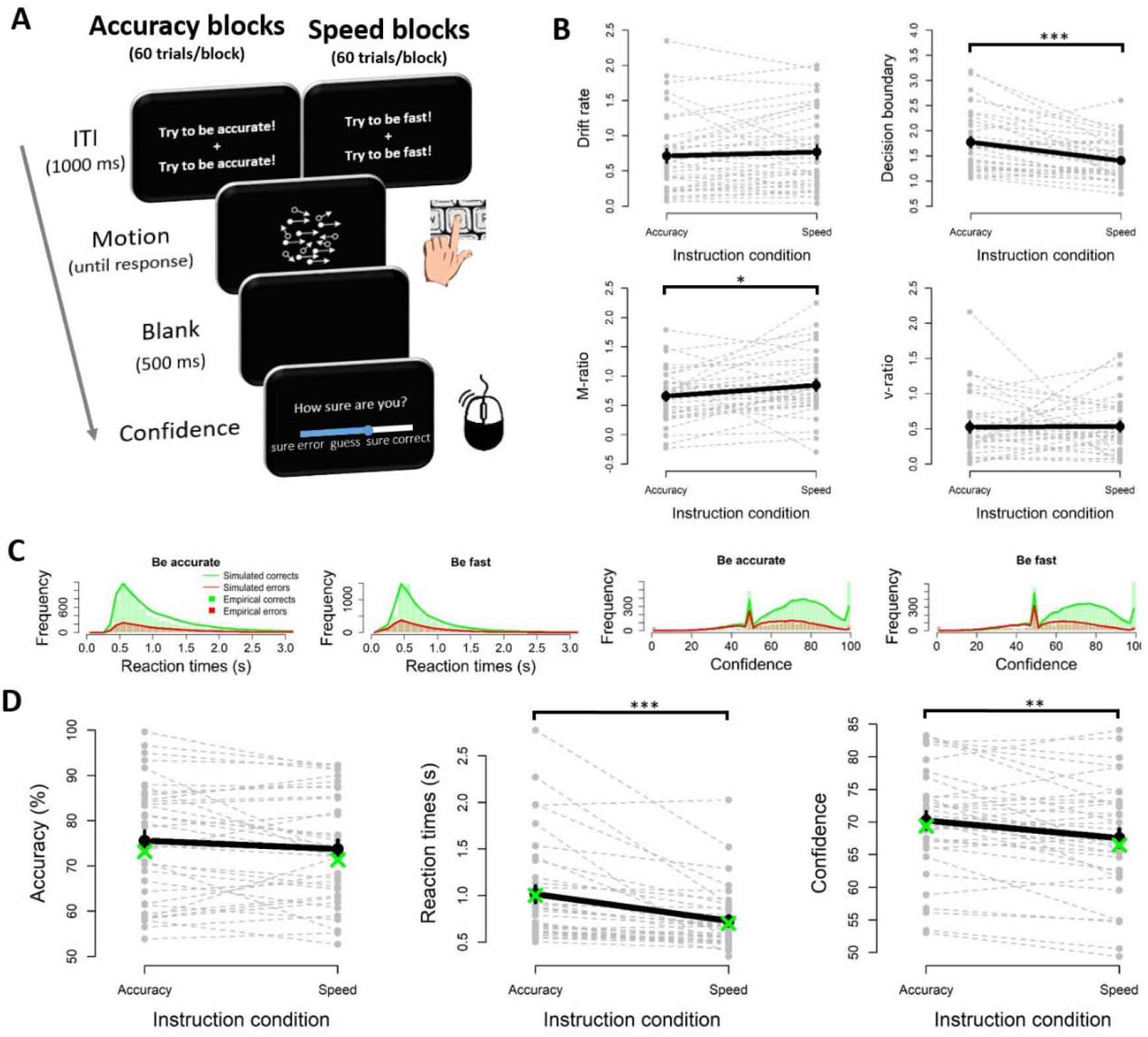
The influence of speed-accuracy instructions on metacognitive accuracy (Experiment 1). **A.** Sequence of events in the experimental task. Participants decided whether the majority of dots were moving left or right, by pressing “E” or “T” with their left hand. After a short blank, they then indicated their level of confidence on a continuous scale. Depending on the block, instructions during the ITI were either to focus on choice accuracy or to focus on speed. **B.** Fitted parameters of a drift diffusion model with additional post-decision accumulation. Fitted decision boundaries were lower in the speed vs accuracy condition, whereas drift rates did not differ. Critically, M-ratio was higher in the speed vs accuracy condition whereas v-ratio did not differ between both instruction conditions. **C.** Distribution of reaction times and confidence for empirical data (bars) and model fits (lines), separately for corrects (green) and errors (red). **D**. Participants were faster, less accurate and less confident when instructed to focus on speed rather than on accuracy. Note: grey lines show individual data points; black lines show averages; green dots show model fits; error bars reflect SEM; ***p<.001, **p<.01, *p<.05.

To shed further light on the underlying cognitive processes, we fitted these data using the evidence accumulation model described in Figure 1A. The basic architecture of our model was a DDM, in which noisy perceptual evidence accumulates over time until a decision boundary is reached. Afterwards, evidence continued to accumulate for a specified amount of time ^27^. In addition to drift rate, decision boundary and non-decision time, our model featured a free parameter controlling the strength of the post-decision evidence accumulation (v-ratio, reflecting the ratio between post-decision drift rate and drift rate) and two further parameters controlling the mapping from *p*(*correct*) onto the confidence scale (see Methods). Generally, our model fitted the data well, as it captured the distributional properties of both reaction times and decision confidence (see Figure 2C). As a first sanity check, we confirmed that decision boundaries were indeed different between the two instruction conditions, *M_speecl_* = 1.40 versus *M_accuracy_* = 1.77, *t*(35) = 4.60, *p* < .001, suggesting that participants changed their decision boundaries as instructed. Also non-decision time tended to be a bit shorter in the speed condition compared to the accuracy condition, *M_speed_* = 309ms versus *Maccdracy* = 390ms, *t*(35) = 3.19, *p* = .003. Drift rates did not differ between both instruction conditions, *p* = .368. There was a small but significant difference between the two instruction conditions in the two additional parameters controlling the idiosyncratic mapping between *p*(*correct*) and the confidence scale, reflecting that in the accuracy condition confidence judgments were slightly higher, *t*(35) = 2.506, *p* = .017, and less variable, *t*(35) = 2.206, *p* = .034, compared to the speed condition.

We next focused on metacognitive accuracy in both conditions (see Figure 2B). In line with the model simulations, our data showed that M-ratio was significantly affected by the speed-accuracy tradeoff instructions, *M_speed_* = 0.84 versus *M_accuracy_* = 0.66, *t*(35) = 2.26, *p* = .030. Moreover, apart from these between-condition differences we also observed significant between-participants correlations between M-ratio and decision boundary both in the accuracy condition, *r*(34) = −.36, *p* = .030, and in the speed condition, *r*(34) = −.53, *p* < .001. Consistent with the notion that metacognitive accuracy should not be affected by differences in decision boundary, v-ratio did not differ between both instruction conditions, *p* = .938. In the model just reported, all parameters were allowed to vary as a function of both instruction conditions. This allowed us to evaluate for each parameter whether or not it is affected by the instruction condition. Note, however, that qualitatively similar results were obtained when instead fixing non-decision time, *M*and *SD* across both conditions.

### Experiment 2: Spontaneous differences in response caution relate to static but not dynamic measures of metacognitive accuracy

Although Experiment 1 provides direct evidence that changes in decision boundary affect M-ratio, it remains unclear to what extent this is also an issue in experiments without speed stress. Notably, in many metacognition experiments, participants do not receive the instruction to respond as fast as possible. Nevertheless, it remains possible that participants implicitly decide on a certain level of response caution. For example, a participant who is eager to finish the experiment quickly might adopt a lower decision boundary compared to a participant who is determined to perform the experiment as accurate as possible, thus leading to natural across-subject variation in decision boundaries. To examine this possibility, in Experiment 2 we analyzed data from an experiment in which participants (N = 63) did not receive any specific instructions concerning speed or choice accuracy. Participants were presented with two boxes filled with dots and had to decide which of the two boxes contained more dots by pressing the “S” or “L” key (corresponding to left and right, respectively). After their choice, participants used the same keys to move a cursor on a continuous confidence scale to indicate their level of confidence (see Figure 3A). The final confidence judgment was confirmed using the enter key. The same evidence accumulation model as before was used to fit these data, and again this model captured both reaction times and decision confidence distributions (Figure 3B). Consistent with our model simulations, model fits showed a positive between-participants correlation between M-ratio and v-ratio, *r*(61) = .21, *p* = .092, although this correlation was not statistically significant (Figure 3C). However, we again observed that M-ratio correlated negatively with the fitted decision boundary, *r*(61) = −.34, *p* = .006, whereas v-ratio did not, *r*(61) = −.19, *p* = .129. To rule out the possibility that the negative relation between decision boundary and M-ratio is confounded by post-decisional processing time, we ran a multiple regression analysis in which we predicted M-ratio by fitted decision boundary and median confidence RTs. As expected, we observed a significant effect of decision boundary, *t*(60) = −2.65, *p* = .010, but not for confidence RTs, |*t*| < .261. Finally, the data showed a significant correlation between median choice RTs and median confidence RTs, *r*(61) = .516, *p* < .001, and between estimated decision boundaries and median confidence RTs, *r*(61) = .427, *p* < .001.

**Figure 3.**
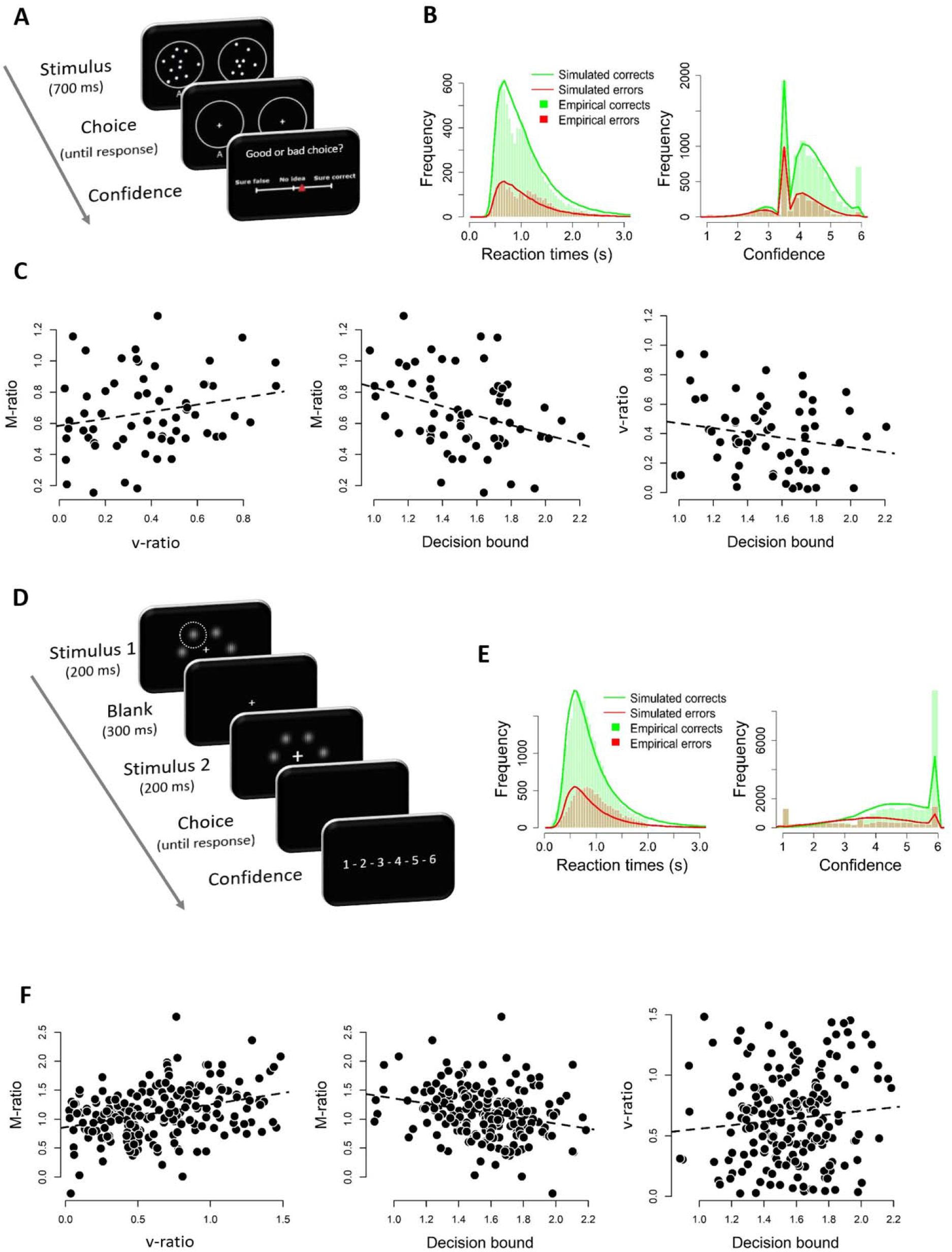
The influence of spontaneous variations in speed-accuracy tradeoff on metacognitive accuracy. **A.** Sequence of events in Experiment 2. On each trial participants decided which of the two circles contained more dots. Afterwards, they indicated their level of confidence on a continuous scale. Note that participants did not receive any instructions concerning speed or accuracy. **B.** Distribution of reaction times and confidence for Experiment 2, using the same conventions as in Figure 2. **C.** The data of Experiment 2 showed a non-significant positive relation between M-ratio and v-ratio (r=.21). Critically, only M-ratio correlated negatively with decision boundary (r=-.34) whereas this relation was not significant for v-ratio (r= −.19). **D.** Sequence of events in Experiment 3. On each trial, participants decided in which temporal interval (first or second) one of the Gabor patches had a higher contrast (highlighted here with a dasher circle which was not present during the experiment). After this choice, participants indicated confidence on a continuous scale. **E.** Distribution of reaction times and confidence for Experiment 3, using the same conventions as in Figure 2. **F.** The data of Experiment 3 showed a significant positive relation between M-ratio and v-ratio (r=.32) and a significant negative correlation between M-ratio and decision boundary (r=-.25) but not between v-ratio and decision boundary (r=.10).

### Experiment 3: Replication in an independent dataset

To assess the robustness of our findings, in Experiment 3 we aimed to replicate our analysis in an independent dataset with high experimental power. To achieve this, we searched the confidence database ^38^ for studies with high power (N > 100) in which a 2CRT task was performed with separate confidence ratings given on a continuous scale. Moreover, because our fitting procedure was not designed for multiple levels of difficulty, we focused on studies with a single difficulty level. We identified one study that satisfied all these constraint (Figure 3D; Prieto, Reyes & Silva, *under review*). In the study by Prieto et al., participants were presented with two consecutive arrays of six Gabor patches, and were asked to decide in which of the two arrays one of the patches had a higher contrast. The response modalities were similar to those used in Experiment 2, but their high experimental power (N=204) assured a very sensitive analysis of our claims. Consistent with the previous analysis, model fits on this independent dataset showed a positive and statistically significant between-participants correlation between M-ratio and v-ratio, *r*(202) = .32, *p* < .001, suggesting that both variables capture shared variance reflecting metacognitive accuracy (see Figure 3F). We again observed that M-ratio correlated negatively with the fitted decision boundary, *r*(202) = −.25, *p* < .001, whereas no relation with decision bound was found for v-ratio, *r*(202) = .10, *p* = .141. A multiple regression analysis further confirmed that variation in M-ratio was explained by variations in decision boundary, *t*(201) = −3.887,*p* < .001, but not in confidence RTs, |*t*| < 1.21, *p* = .225. Finally, the data showed a significant correlation between median choice RTs and median confidence RTs, *r*(202) = .277, *p* < .001, and between estimated decision boundaries and median confidence RTs, *r*(202) = .151, *p* = .031.

### Relating decision boundary to *d*’ and meta-*d*’

Given that M-ratio reflects the ratio between *d*’ and meta-*d*’ it is instructive to further unravel the relation of both these measures with the decision boundary. Interestingly, whereas we observed a clear negative relation between M-ratio and decision boundary in both the model simulations and all three experiments, the findings concerning *d*’ and meta-*d*’ are much less straightforward. In the simulations, we observed a significant positive correlation between *d*’ and decision boundary, *r*(98) = .589,*p* < .001, but not between meta-*d*’ and decision boundary, *r*(98) = .155, *p* = .124. In Experiment 1, although M-ratio was modulated by instruction condition, this was not the case for *d*’, *p* = .118, or meta-*d*’, *p* = .300. In Experiment 2, we observed a negative relation between decision boundary and *d*’, *r*(61) = −.306,*p* = .015, and a negative relation between decision boundary and meta-*d*’, *r*(61) = −.423, *p* < .001. Finally, in Experiment 3, we observed a positive relation between decision boundary and *d*’, *r*(202) = .250,*p* < .001, and a non-significant negative relation between decision boundary and meta-*d*’, *r*(202) = −.097, *p* = .167.

## Discussion

Researchers across several fields show an increasing interest in the question how observers can evaluate their task performance via confidence judgments. Crucial to study metacognition is a method to objectively quantify the extent to which participants are able to detect their own mistakes, regardless of decision strategy. We here report that a commonly used *static* measure of metacognitive accuracy (M-ratio) highly depends on the decision boundary – reflecting response caution – that is set for decision making. This was the case in simulation results, in an experiment explicitly manipulating the tradeoff between speed and accuracy, and in two datasets in which participants received no specific instructions concerning speed or accuracy. We propose an alternative, *dynamic*, measure of metacognitive accuracy (v-ratio) that does not depend on decision boundary.

### Caution is warranted with static measures of metacognition

The most important consequence of the current findings is that researchers should be cautious when interpreting static measures of metacognitive accuracy, such as M-ratio. Although the findings reported in Experiments 2 and 3 are correlational and should thus be interpreted with caution (e.g., it could be that participants with good metacognition deem it appropriate to impose low decision boundaries), the reported simulations and the within-participant experimental manipulation of Experiment 1 are indicative of a fundamental issue with M-ratio. Moreover, differences in confidence between the two instruction conditions in Experiment 1 were rather subtle, suggesting that even minor influences in decision confidence are sufficient to induce differences in M-ratio. As the name indicates, M-ratio reflects the ratio between meta-*d*’ (second-order performance) and *d*’ (first-order performance). Interestingly, whereas both the simulations and the three experiments showed consistent associations between M-ratio and decision boundary, the story was more complicated when instead directly relating meta-*d*’ and *d*’ with variations in decision boundary. Whereas in the simulations the relation between M-ratio and decision boundary was largely driven by *d*’ but not so much by meta-*d*’, these results were less clear in the empirical experiments.

In the following, we will discuss several examples where our finding might have important implications. In the last decade there has been quite some work investigating to what extent the metacognitive evaluation of choices is a domain-general process or not. These studies often require participants to perform different kinds of tasks, and then examine correlations in choice accuracy and in metacognitive accuracy between these tasks ^3,17–20,39^. For example, Mazancieux and colleagues asked participants to perform an episodic memory task, a semantic memory task, a visual perception task and a working memory task. In each task, participants rated their level of confidence after a decision. The results showed that whereas correlations between choice accuracy on these different tasks were limited, there was substantial covariance in metacognitive accuracy across these domains. Because in this study participants received no time limit to respond, it remains unclear whether this finding can be interpreted as evidence for a domain-general metacognitive monitor, or instead a domain-general response caution which caused these measures to correlate. Another popular area of investigation has been to unravel the neural signatures supporting metacognitive accuracy ^19,20,40–42^. For example, McCurdy et al. observed that both visual and memory metacognitive accuracy correlated with precuneus volume, potentially pointing towards a role of precuneus in both types of metacognition. It remains unclear, however, to what extent differences in response caution might be responsible for this association. Although differences in response caution are usually found to be related to pre-SMA and anterior cingulate ^24,25^, there is some suggestive evidence linking precuneus to response caution ^43^. Therefore, it is important that future studies on neural correlates of metacognition rule out the possibility that their findings are caused by response caution. Finally, our study has important consequences for investigations into differences in metacognitive accuracy between specific, e.g. clinical, groups. For example, Folke and colleagues ^23^ reported that M-ratio was reduced in a group of bilinguals compared to a matched group of monolinguals. Interestingly, they also observed that on average bilinguals had shorter reaction times than monolinguals, but this effect was unrelated to the group difference in M-ratio. Because these authors did not formally model their data using evidence accumulation models, however, it remains unclear whether this RT difference results from a difference in boundary, and if so to what extent this explains the difference in M-ratio between both groups that was observed. In a similar vein, individual differences in M-ratio have been linked to psychiatric symptom dimensions, and more specifically to a symptom dimension related to depression and anxiety ^5^. At the same time, it is also known that individual differences in response caution are related to a personality trait known as *need for closure* ^44^ Given that need for closure is, in turn, related to anxiety and depression ^45^, it remains a possibility that M-ratio is only indirectly related to these psychiatric symptoms via response caution.

### The potential of dynamic measures of metacognition

In order to control for potential influences of response caution on measures of metacognitive accuracy, one approach could be to estimate the decision boundary and examine whether the relation between metacognitive accuracy and the variable of interest remains when controlling for decision boundary (e.g., using mediation analysis). However, a more direct approach would be to estimate metacognitive accuracy in a dynamic framework, thus taking into account differences in response caution. For example, building on the drift diffusion model it has been proposed that confidence reflects the probability of a choice being correct given evidence, decision time, and the decision that was made ^32–34^. Usually, it is (implicitly) assumed that participants learn the mapping between these variables via experience ^32,46^, however the precise learning mechanism remains to be unraveled. In the current work, we proposed v-ratio (reflecting the ratio between post-decision drift rate and drift rate) as such a dynamic measure of metacognitive accuracy (following the observation that post-decision drift rate indexes how accurate confidence judgments are^27,28^). In both simulations and empirical data, we observed a positive relation between v-ratio and M-ratio, suggesting they capture shared variance. Critically, v-ratio was not correlated with decision boundary, suggesting it is not affected by differences in response caution. Thus, our dynamic measure of metacognition holds promise as a novel approach to quantify metacognitive accuracy.

In our approach we allowed the drift rate and the post-decision drift rate to dissociate. This proposal is in line with the view of metacognition as a second-order process whereby dissociations between confidence and choice accuracy might arise because of noise or bias at each level ^47–49^. However, when formulating post-decision drift rate as a continuation of evidence accumulation, it remains underspecified which evidence the post-decision accumulation process is exactly based on. It has been suggested that participants can accumulate evidence that was still in the processing pipeline (e.g. in a sensory buffer) even after a choice was made ^36,50^. However, it is not very likely that this is the only explanation, particularly in tasks without much speed stress. One other likely possibility, is that during the post-decision process, participants resample the stimulus from short-term memory ^51^. Because memory is subject to decay, dissociations between the post-decision drift rate and the drift rate can arise. Other sources of discrepancy might be contradictory information quickly dissipating from memory ^52^ which should lower metacognitive accuracy, or better assessment of encoding strength with more time ^53^ which should increase metacognitive accuracy.

One important caution is that in our proposed formalization the computation of decision confidence (and thus metacognitive accuracy) arises by means of post-decisional evidence accumulation. Put simply, an observer with post-decision drift rate of one will be good at telling apart corrects from errors whereas an observer with post-decision drift rate of zero will be at chance level. One way to evaluate the validity of post-decisional drift rate for this purpose, is to simulate data with various levels of post-decision drift rate, and then evaluate whether the area under the Type-II ROC is different from chance level. Type-II ROC analysis is a bias free measure that quantifies how good confidence tracks accuracy^16^. As shown in Supplementary Note 2, this analysis indeed reveals a chance level area under type-II ROC whenever post-decision drift rate equals zero. However, this is only the case when a single level of difficulty is used (as was the case in all experiments reported here). When multiple levels of difficulty are used, this is no longer the case. Although all datasets reported in the current study feature only a single level of difficulty, and thus our model can be validly used, we nevertheless fitted our data with a variant of the model in which confidence is determined by the level of post-decisional evidence (following ^27,28^), rather than by (post-decisional) probability of success, as in our original formulation. The latter formulation of the dynamic model has the advantage that it shows chance level area under type II ROC when post-decision drift rate equals zero even when there are multiple levels of difficulty. Most importantly, the findings described in the current manuscript are not contingent upon the specific implementation of confidence. As shown in Supplementary Note 3, both variants of the dynamic model provide highly similar results. Therefore, we deposited code for both versions of the model so that researchers interested in computing v-ratio can fit the model best suited for their data. Finally, it is important to stress that rather than arbitrating between two highly similar models, the main goal of the current work was to demonstrate that in order to correctly quantify metacognitive accuracy it is critical to rely on a dynamic model, such as the two variants that we report in the current paper.

Finally, we note that in the modeling efforts reported here, the duration of post-decisional evidence accumulation was simply taken to be the median of confidence RTs. Most previous modeling efforts have likewise assumed that post-decision processing terminates once confidence is externally queried^28^, and only a few studies have explicitly examined different stopping rules for post-decision processing ^46,54^. Given that we still lack a clear mechanistic understanding of how post-decisional processing is terminated, we here decided for this pragmatic implementation. However, by further unravelling the computational mechanisms underlying post-decisional accumulation termination, substantial progress can still be made by including these mechanisms in future modeling efforts. To sum up, we provided evidence from simulations and empirical data that a common static measure of metacognition, M-ratio, is confounded with response caution. We proposed an alternative measure of metacognition based on a dynamic framework, v-ratio, which is insensitive to variations in caution, and may thus be suitable to study how metacognitive accuracy varies across subjects and conditions.

## Methods

### Computational model

#### Simulations

Data were simulated for 100 observers with 1000 trials each. Note that the findings concerning M-ratio and decision boundary remain qualitatively unchanged when instead simulating more observers (500), more trials per observer (5000) or both. For each simulated observer, we randomly selected a value for the drift rate (uniform distribution between 0 and 2.5), for the decision boundary (uniform distribution between .5 and 3), for the non-decision time (uniform distribution between .2 and .6) and for the v-ratio (uniform distribution between 0 and 1.5; see below for details). When v-ratio is above 1, this implies that observers use more information evaluate their choices then they did to make the actual choice^55^. To estimate meta-*d*’, data is needed for both of the possible stimuli (i.e., to estimate bias); therefore, for half of the trials we multiplied the drift rate by −1. During model fitting, we used empirically observed confidence RTs to inform the duration of post-decision processing time (see below). Given that the empirical data showed moderate correlations between choice RTs and confidence RTs (i.e., .277 and .516 for Experiment 2 and 3), for the model simulations we generated post-decision processing times that were moderately correlated with the choice RTs. To achieve this, for each simulated observer we first generated dummy data using the relevant combination of bound, drift and non-decision time, and during the actual simulations post-decision processing time was sampled from a normal distribution (sigma = 1) around the mean of correct choice RTs in the dummy data. This procedure induced a moderate correlation between choice RTs and confidence RTs, *r*(98) = 32, *p* < .001, and between confidence RTs and decision boundary, *r*(98) = .32, *p* = .001. Finally, we fixed the values for starting point *(z* = .5), within-trial noise (*σ* = 1).

#### Fitting procedure

We coded an extension of the drift diffusion model (DDM) that simultaneously fitted choices, reaction times and decision confidence. The standard DDM is a popular variant of sequential sampling models of two-choice tasks. We used a random walk approximation, coded in the rcpp R package to increase speed ^56^, in which we assumed that noisy sensory evidence started at *z*a;* 0 and a are the lower and upper boundaries, respectively, and *z* quantifies bias in the starting point (z = .5 means no bias). At each time interval *τ* a displacement Δ in the integrated evidence occurred according to the formula shown in equation (1):

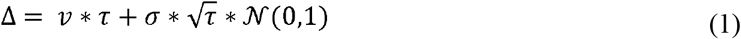

Evidence accumulation strength is controlled by *v*, representing the drift rate, and within-trial variability, σ, was fixed to 1. Note that it is common practice in DDM fitting to fix within-trial variability to 1, although this assumption of constant within-trial noise is often not made explicitly. The reason for fixing this parameter is that changes in σ cannot be dissociated from changes in *a*. Given that our hypothesis specifically concerns the latter, we decided to fix σ to 1. The random walk process continued until the accumulated evidence crossed either 0 or *a*. After boundary crossing, the evidence continued to accumulate for a duration depending on the participant-specific median confidence reaction time. Specifically, for each participant we calculated the difference in time between the moment that participants made their initial choice and the moment that they confirmed their confidence judgment. From these differences we calculated, per participant, the median and used this value as the duration of post-decision processing time. The rationale for this choice was that it reflects a good approximation of the time participants effectively continued to accumulate additional evidence following an initial choice. Note that during the post-decisional accumulation period the integrated evidence was not limited between the boundaries (i.e. 0 was not a hard boundary) and could land anywhere between ∞ and +∞o. Importantly, consistent with the signal detection theoretical notion that primary and secondary evidence can dissociate, we allowed for dissociations between the drift rate governing the choice and the post-decision drift rate. For compatibility with the M-ratio framework, we quantified metacognitive accuracy as the ratio between post-decision drift rate and drift rate, as shown in equation (2):

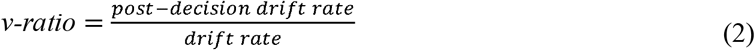

As a consequence, when v-ratio = 1, this implies that post-decision drift and drift are the same. When v-ratio = .5, the magnitude of the post-decision drift rate is half the magnitude of the drift rate. To calculate decision confidence, we first quantified for each trial the probability of being correct given evidence, time, and choice. The heat map representing *p*(*correct/e, t, X*) is shown in Figure 1A, and was constructed by means of 300.000 random walks without absorbing bounds, with drift rates sampled from a uniform distribution between zero and ten. This assured sufficient data points across the relevant part of the heat map. Subsequently, the average accuracy was calculated for each (response time, evidence, choice) combination, based on all trials that had a data point for that (response time, evidence, choice) combination. Smoothing was achieved by aggregating over evidence windows of .01 and *τ* windows of 3. Next, to take into account idiosyncratic mappings of *p*(*correct/e, t, X*) onto the confidence scale used in the experiment, we added two extra free parameters that controlled the mean (M) and the width (SD) of confidence estimates, as shown in equation (3):

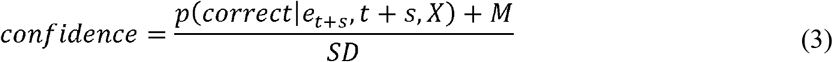

Although empirical confidence distributions appeared approximately normally distributed, there was an over-representation of confidence values at the boundaries (1 and 100 in Experiment 1; 1 and 6 in Experiments 2 and 3) and in the middle of the scale (50 in Experiment 1, 3.5 in Experiment 2). Most likely, this resulted from the use of verbal labels placed at exactly these values. To account for frequency peaks at the endpoints of the scale, we relabeled predicted confidence values that exceeded the endpoints of the scale as the corresponding endpoint (e.g., in Experiment 1 a predicted confidence value of 120 was relabeled as 100), which naturally accounted for the frequency peaks at the endpoints. To account for peaks in the center of the scale, we assumed that confidence ratings around the center were pulled towards the center value. Specifically, we relabeled *P*% of trials around the midpoint as the midpoint (e.g., in Experiment 1, *P* = 10% implies that 10% of the data closest to 50 were (re)labeled as 50). Note that *P* was not a free parameter, but instead its value was taken to be the participant-specific proportion based on the empirical data. Note that the main conclusions reported in this manuscript concerning the relation between M-ratio, decision boundary and post-decision drift rate, remain the same in a reduced model without *P*, and also in a reduced model without *P, M* and *SD*. Because these reduced models did not capture confidence distributions very well though, we here report only the findings of the full model.

To estimate these 6 parameters (*v, a, Ter, v-ratio, M, and SD*) based on choices, reaction times and decision confidence, we implemented quantile optimization. Specifically, we computed the proportion of trials in quantiles .1, .3, .5, .7, and .9, for both reaction times and confidence; separately for corrects and errors (maintaining the probability mass of corrects and errors, respectively). We then used differential evolution optimization, as implemented in the DEoptim R package ^57^, to estimate these 6 parameters by minimizing the chi square error function shown in equation (4):

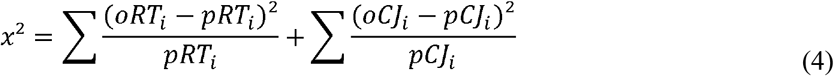

with oRT_i_ and pRT_i_ corresponding to the proportion of observed/predicted responses in quantile *i*, separately calculated for corrects and errors both reaction times, and oCJ_i_ and pCJ_i_ reflecting their counterparts for confidence judgments. We set *τ* to .001. Model fitting was done separately for each participant. Note that in Experiment 3 there was no clear peak in the middle of the scale so *P* was fixed to 0 in that experiment.

M-ratio, reflecting the ratio between meta-*d*’ and *d*’, was calculated using the method described in Maniscalco and Lau ^15^. Because confidence was expressed on a continuous scale in the simulations and all three experiments, before fitting the meta-*d*’ model (which requires categorical confidence judgments) confidence was divided into 4 bins, separately for each participant, before fitting the meta-*d*’ model.

#### Parameter recovery

To assure that our model was able to recover the parameters, we here report parameter recovery. In order to assess parameter recovery with a sensible set of parameter combinations, we used the fitted parameters of Experiment 1 (N = 36), simulated data from these parameters with a varying number of trials, and then tested whether our model could recover these initial parameters. As a sanity check, we first simulated a large number of trials (25000 trials per participant), which as expected provided excellent recovery for all six parameters, *r*s > .97. We then repeated this process with only 200 trials per participants, which was the trial count in Experiment 2 (note that Experiment 1 and 3 both had higher trial counts). Recovery for v-ratio was still very good, *r* = .85, whereas it remained excellent for all other parameters, *r*s > .98.

### Experiment 1

#### Participants

Forty healthy participants (18 males) took part in Experiment 1 in return for course credit (mean age = 19.82, between 18 and 30). All reported normal or corrected-to-normal vision. Two participants were excluded because they required more than 10 practice blocks in one of the training blocks (see below) and two participants were excluded because their choice accuracy, averaged per block and then compared against chance level using a one-sample t-test, was not significantly above chance level. The final sample thus comprised thirty-six participants. All participants provided their informed consent and all procedures adhered to the general ethical protocol of the ethics committee of the Faculty of Psychology and Educational Sciences of Ghent University.

#### Stimuli and apparatus

The data for Experiment 1 were collected in an online study, due to COVID-19. As a consequence, caution is warranted when interpreting between-participant differences given that we had no control over hardware specifications. Given that Experiment 1 concerns a within-subject experimental manipulation, however, the risk that this is problematic for our data is small. Participants were allowed to take part in the experiment only when they made us of an external mouse. Choices were provided with the keyboard, and decision confidence was indicated with the mouse. Stimuli in Experiment 1 consisted of 50 randomly moving white dots (radius: 2 pixels) drawn in a circular aperture on a black background centered on the fixation point. Dots disappeared and reappeared every 5 frames. The speed of dot movement (number of pixel lengths the dot will move in each frame) was a function of the screen resolution (screen width in pixels / 650).

#### Task procedure

Each trial started with the presentation of a fixation cross for 1000 ms. Above and below this fixation cross specific instructions were provided concerning the required strategy. In accuracy blocks the instruction was to respond as accurately as possible; in speed blocks the instruction was to respond as fast as possible. The order of this block-wise manipulation was counterbalanced across participants. Next, randomly moving dots were shown on the screen until a response was made or the response deadline was reached (max 5000 ms). On each trial, 20% of the dots coherently moved towards the left or the right side of the screen, with an equal number of leftward and rightward movement trials in each block. Participants were instructed to decide whether the majority of dots was moving towards the left or the right side of the screen, by pressing “E” or “T”, respectively, with their left hand. After their response, a blank screen was shown for 500 ms, followed by the presentation of a continuous confidence scale. Below the scale the labels “Sure error”, “guess”, and “sure correct” were shown, arranged outer left, centrally and outer right, respectively. After clicking the confidence scale, participants had to click a centrally presented “Continue” button (below the confidence scale) that ensured that the position of the mouse was central and the same on each trial.

The main part of Experiment 1 consisted of 10 blocks of 60 trials, half of which were from the accuracy instruction condition and half from the speed instruction condition. The experiment started with 24 practice trials during which participants only discriminated random dot motion at 50% coherence, no confidence judgments were asked. This block was repeated until participants achieved 85% accuracy (mean = 2 blocks). Next, participants completed again 24 practice trials with the only difference that now the coherence was decreased to 20% (mean = 1.05 blocks). When participants achieved 60% accuracy, they then performed a final training block of 24 trials during which they practiced both dot discrimination and indicated their level of confidence (mean = 1.05 blocks).

### Experiment 2

Full experimental details are described in Drescher et al. ^58^. Participants were seated in individual cubicles in front of a 15-in CRT monitor with a vertical refresh rate of 85Hz. On each trial participants were presented with two white circles (5.1° diameter) on a black background, horizontally next to each other with a distance of 17.8° between the midpoints. Fixation crosses were shown for 1s in each circle, followed by dots clouds in each circle for 700ms. The dots had a diameter of 0.4°. Dot positions in the boxes, as well as the position of the box containing more dots were randomly selected on each trial. The difference in number of dots between both boxes (indexing task difficulty) was adapted online using an unequal step size staircase procedure^20^. Participants indicated which circle contained more dots by pressing “S” or “L” on a keyboard. Then, the question “correct or false?” appeared on the screen, with a continuous confidence rating bar, with the labels “Sure false”, “No idea”, and “Sure correct”. Participants moved a cursor with the same keys as before, and confirmed their confidence judgment with the enter key. No time limit was imposed for both primary choice and confidence rating. Subjects received several practice trials (10 without confidence rating, 14 with confidence rating), before they completed eight experimental blocks of 25 trials.

### Experiment 3

Data from this experiment were taken from the confidence database ^59^, a collection of openly available studies on decision confidence. In this experiment, Prieto, Reyes and Silva ^60^, used the same task as described in Fleming and colleagues ^2^. Each participant (N=204 woman, aged 18-35) completed 50 practice trials, followed by 5 blocks of 200 trials. On each trial participants were presented with an array of eight Gabor patches for 200ms, a blank for 300ms and another array of 6 Gabor patches for 200ms. Participants had to decide whether the first or the second temporal interval contained a patch with a higher contrast. The contrast of the pop-out Gabor was continuously adapted using an online staircase procedure to maintain 71% accuracy. Choices and confidence reports were collected in an identical manner as in Experiment 2. The only difference was that there was no verbal description around the middle point of the confidence scale.

## Data and code availability

All experiment code, analysis code and raw data have been deposited online and can be freely accessed (github.com/kdesende/dynamic_influences_on_static_measures/).

## Acknowledgments

The authors like to thank Peter R Murphy, Bharath Chandra Talluri, and Annika Boldt for insightful discussions and Pierre Le Denmat, Robin Vloeberghs and Alan Voodla for comments on an earlier draft. This research was supported by an FWO [PEGASUS]^2^ Marie Skłodowska-Curie fellowship (12T9717N, to K.D.) and a starting grant from the KU Leuven (STG/20/006, to K.D.). L.V. was supported by the FWO (11H5619N)

## References

1. Sanders, J. I., Hangya, B. & Kepecs, A. Signatures of a Statistical Computation in the Human Sense of Confidence. Neuron 90, 499–506 (2016).

2. Fleming, S. M., Weil, R. S., Nagy, Z., Dolan, R. J. & Rees, G. Relating introspective accuracy to individual differences in brain structure. Science (80-.). 329, 1541–3 (2010).

3. Baird, B., Mrazek, M. D., Phillips, D. T. & Schooler, J. W. Domain-Specific Enhancement of Metacognitive Ability Following Meditation Training. J. Exp. Psychol. Gen. 143, 1972–1979 (2014).

4. Rollwage, M. et al. Metacognitive failure as a feature of those holding radical political beliefs. Nat. Hum. Behav. 2, 637–644 (2018).

5. Rouault, M., Seow, T., Gillan, C. M. & Fleming, S. M. Psychiatric Symptom Dimensions Are Associated With Dissociable Shifts in Metacognition but Not Task Performance. Biol. Psychiatry 84, 443–451 (2018).

6. Locke, S. M., Mamassian, P. & Landy, M. S. Performance monitoring for sensorimotor confidence: A visuomotor tracking study. Cognition 205, (2020).

7. Arbuzova, P. et al. Measuring Metacognition of Direct and Indirect Parameters of Voluntary Movement Polina. bioxRxiv (2020).

8. Yallak, E. & Balci, F. Metric error monitoring: Another generalized mechanism for magnitude representations? Cognition 210, 104532 (2021).

9. De Martino, B., Fleming, S. M., Garrett, N. & Dolan, R. J. Confidence in value-based choice. Nat. Neurosci. 16, 105–10 (2013).

10. Bang, D. et al. Confidence matching in group decision-making. Nat. Hum. Behav. 1, 1–7 (2017).

11. Meyniel, F., Schlunegger, D. & Dehaene, S. The Sense of Confidence during Probabilistic Learning: A Normative Account. PLoS Comput. Biol. 11, e1004305 (2015).

12. Nelson, T. O. A comparison of current measures of the accuracy of feeling-of-knowing predictions. Psychol. Bull. 95, 109–133 (1984).

13. Sandberg, K., Timmermans, B., Overgaard, M. & Cleeremans, A. Measuring consciousness: is one measure better than the other? Conscious. Cogn. 19, 1069–78 (2010).

14. Fleming, S. M. HMeta-d: hierarchical Bayesian estimation of metacognitive efficiency from confidence ratings. Neurosci. Conscious. 3, 1–14 (2017).

15. Maniscalco, B. & Lau, H. A signal detection theoretic approach for estimating metacognitive sensitivity from confidence ratings. Conscious. Cogn. 21, 422–30 (2012).

16. Fleming, S. M. & Lau, H. C. How to measure metacognition. Front. Hum. Neurosci. 8, 1–9 (2014).

17. Mazancieux, A. et al. Is There a G Factor for Metacognition ? Correlations in Retrospective Metacognitive Sensitivity Across Tasks Is There a G Factor for Metacognition ? Correlations in Retrospective Metacognitive Sensitivity Acros. J. Exp. Psychol. Gen. (2020).

18. Rouault, M., McWilliams, A., Allen, M. G. & Fleming, S. M. Human Metacognition Across Domains: Insights from Individual Differences and Neuroimaging. Personal. Neurosci. 1, (2018).

19. McCurdy, L. Y. et al. Anatomical coupling between distinct metacognitive systems for memory and visual perception. J. Neurosci. 33, 1897–906 (2013).

20. Fleming, S. M., Ryu, J., Golfinos, J. G. & Blackmon, K. E. Domain-specific impairment in metacognitive accuracy following anterior prefrontal lesions. Brain 137, 2811–2822 (2014).

21. Filevich, E., Dresler, M., Brick, X. T. R. & Ku, S. Metacognitive Mechanisms Underlying Lucid Dreaming. J. Neurosci. 35, 1082–1088 (2015).

22. Rahnev, D. a, Maniscalco, B., Luber, B., Lau, H. C. & Lisanby, S. H. Direct injection of noise to the visual cortex decreases accuracy but increases decision confidence. J. Neurophysiol. 107, 1556–1563 (2012).

23. Folke, T., Ouzia, J., Bright, P., De Martino, B. & Filippi, R. A bilingual disadvantage in metacognitive processing. 150, 119–132 (2016).

24. Forstmann, B.U. et al. Striatum and pre-SMA facilitate decision-making under time pressure. Proc. Natl. Acad. Sci. U. S. A. 105, 17538–42 (2008).

25. Bogacz, R., Wagenmakers, E.-J., Forstmann, B. U. & Nieuwenhuis, S. The neural basis of the speed-accuracy tradeoff. Trends Neurosci. 33, 10–6 (2010).

26. Scheffers, M. K. & Coles, M. G. Performance monitoring in a confusing world: error-related brain activity, judgments of response accuracy, and types of errors. J. Exp. Psychol. Hum. Percept. Perform. 26, 141–151 (2000).

27. Pleskac, T. J. & Busemeyer, J. R. Two-stage dynamic signal detection: A theory of choice, decision time, and confidence. Psychol. Rev. 117, 864–901 (2010).

28. Yu, S., Pleskac, T. J. & Zeigenfuse, M. D. Dynamics of Post-decisional Processing of Confidence. J. Exp. Psychol. Gen. 144, 489–510 (2015).

29. Gold, J. I. & Shadlen, M. N. The neural basis of decision making. Annu. Rev. Neurosci. 30, 535–561 (2007).

30. Ratcliff, R. & McKoon, G. The Diffusion Decision Model□: Theory and Data for Two-Choice Decision Tasks. Neural Comput. 20, 873–922 (2008).

31. Forstmann, B. U. & Wagenmakers, E.-J. Sequential Sampling Models in Cognitive Neuroscience: Advantages, Applications, and Extensions. Annu. Rev. Psychol. 67, 641–666 (2016).

32. Kiani, R., Corthell, L. & Shadlen, M. N. Choice Certainty Is Informed by Both Evidence and Decision Time. Neuron 84, 1329–1342 (2014).

33. Zylberberg, A., Fetsch, C. R. & Shadlen, M. N. The influence of evidence volatility on choice, reaction time and confidence in a perceptual decision. Elife 5, 1–31 (2016).

34. Drugowitsch, J., Moreno-Bote, R., Churchland, A. K., Shadlen, M. N. & Pouget, A. The Cost of Accumulating Evidence in Perceptual Decision Making. J. Neurosci. 32, 3612–3628 (2012).

35. Moreno-Bote, R. Decision confidence and uncertainty in diffusion models with partially correlated neuronal integrators. Neural Comput. 22, 1786–1811 (2010).

36. Van Den Berg, R. et al. A common mechanism underlies changes of mind about decisions and confidence. Elife 1–21 (2016) doi:10.7554/eLife.12192.

37. Desender, K., Donner, T. H. & Verguts, T. Dynamic expressions of confidence within an evidence accumulation framework. Cognition (2020).

38. Rahnev, D. et al. The Confidence Database. Nat. Hum. Behav. 4, (2020).

39. Song, C. et al. Relating inter-individual differences in metacognitive performance on different perceptual tasks. Conscious. Cogn. 20, 1787–92 (2011).

40. Bor, D., Schwartzman, D. J., Barrett, A. B. & Seth, A. K. Theta-burst transcranial magnetic stimulation to the prefrontal or parietal cortex does not impair metacognitive visual awareness. PLoS One 1, 165–175 (2016).

41. Rounis, E., Maniscalco, B., Rothwell, J. J. C., Passingham, R. R. E. & Lau, H. H. Theta-burst transcranial magnetic stimulation to the prefrontal cortex impairs metacognitive visual awareness. Cogn. Neurosci. 1, 165–175 (2010).

42. Baird, B., Cieslak, M., Smallwood, J., Grafton, S. T. & Schooler, J. W. Regional White Matter Variation Associated with Domain-specific Metacognitive Accuracy. J. Cogn. Neurosci. 1–10 (2014) doi:10.1162/jocn.

43. Van Maanen, L. et al. Neural Correlates of Trial-to-Trial Fluctuations in Response Caution. J. Neurosci. 31, 17488–17495 (2011).

44. Evans, N. J., Rae, B., Bushmakin, M., Rubin, M. & Brown, S. D. Need for closure is associated with urgency in perceptual decision-making. Mem. Cogn. 45, 1193–1205 (2017).

45. Freeman, D. et al. Delusions and decision-making style: Use of the Need for Closure Scale. Behav. Res. Ther. 44, 1147–1158 (2006).

46. Desender, K., Donner, T. & Verguts, T. Dynamic expressions of confidence within an evidence accumulation framework. Cognition 104522 (2020) doi:10.1101/2020.02.18.953778.

47. Fleming, S. M. & Daw, N. D. Self-evaluation of decision performance: A general Bayesian framework for metacognitive computation. Psychol. Rev. 124, 1–59 (2016).

48. Pasquali, A., Timmermans, B. & Cleeremans, A. Know thyself: metacognitive networks and measures of consciousness. Cognition 117, 182–90 (2010).

49. Balsdon, T., Wyart, V. & Mamassian, P. Confidence controls perceptual evidence accumulation. Nat. Commun. 11, (2020).

50. Resulaj, A., Kiani, R., Wolpert, D. M. & Shadlen, M. N. Changes of mind in decision-making. Nature 461, 263–266 (2009).

51. Vlassova, A. & Pearson, J. Look Before You Leap: Sensory Memory Improves Decision Making. Psychol. Sci. 24, 1635–1643 (2013).

52. Minson, J. A. & Umphres, C. Confidence in Context: Perceived Accuracy of Quantitative Estimates Decreases With Repeated Trials. Psychol. Sci. (2020) doi:10.1177/0956797620921517.

53. Nelson, T. O. & Dunlosky, J. When People’s Judgments of Learning are extremely accurate at predicting subsequent recall: The ‘Delayed-JOL effect’. Psychol. Sci. 2, 267–270 (1991).

54. Moran, R., Teodorescu, A. R. & Usher, M. Post choice information integration as a causal determinant of confidence: Novel data and a computational account. Cogn. Psychol. 78, 99–147 (2015).

55. Moreira, C. M., Rollwage, M., Kaduk, K., Wilke, M. & Kagan, I. Post-decision wagering after perceptual judgments reveals bi-directional certainty readouts. Cognition 176, 40–52 (2018).

56. Eddelbuettel, D. Seamless R and C++ integration with Rcpp. Seamless R C+ + Integr. with Rcpp 40, 1–220 (2013).

57. Mullen, K. M., Ardia, D., Gil, D. L., Windover, D. & Cline, J. DEoptim: An R package for global optimization by differential evolution. J. Stat. Softw. 40, 1–26 (2011).

58. Drescher, L. H., Van den Bussche, E. & Desender, K. Absence without leave or leave without absence: Examining the interrelations among mind wandering, metacognition, and cognitive control. PLoS One (2018).

59. Rahnev, D. et al. The Confidence Database. Nat. Hum. Behav. (2020) doi:10.1038/s41562-019-0813-1.

60. Prieto, F., Reyes, G. & Silva, J. Role of Maternal Metacognition and Maternal Mental Health in Caregiving Behavior. (2020).

